# Same law, diverging practice: comparative analysis of Endangered Species Act consultations by two federal agencies

**DOI:** 10.1101/165647

**Authors:** Megan Evansen, Ya-Wei Li, Jacob Malcom

## Abstract

Evaluating how wildlife conservation laws are implemented is critical for safeguarding biodiversity. Two agencies, the U.S. Fish and Wildlife Service and National Marine Fisheries Service (FWS and NMFS; Services collectively), are responsible for implementing the U.S. Endangered Species Act (ESA), which requires federal protection for threatened and endangered species. FWS and NMFS’ comparable role for terrestrial and marine taxa, respectively, provides the opportunity to examine how implementation of the same law varies between agencies. We analyzed how the Services implement a core component of the ESA, section 7 consultations, by objectively assessing the contents of >120 consultations on sea turtle species against the requirements in the Services’ consultation handbook, supplemented with in-person observations from Service biologists. Our results showed that NMFS consultations were 1.40 times as likely to have higher completeness scores than FWS consultations given the standard in the handbook. Consultations tiered from an FWS programmatic consultation inherited higher quality scores of generally more thorough programmatic consultations, indicating that programmatic consultations could increase the quality of consultations while improving efficiency. Both agencies commonly neglected to account for the effects of previous consultations and the potential for compounded effects on species. From these results, we recommend actions that can improve quality of consultation, including the use of a single database to track and integrate previously authorized harm in new analyses and the careful but more widespread use of programmatic consultations. Our study reveals several critical shortfalls in the current process of conducting ESA section 7 consultations that the Services could address to better safeguard North America’s most imperiled species.

## 1. INTRODUCTION

The U.S. Endangered Species Act (ESA) is considered one of the strongest wildlife laws in the world (1). Signed into law in 1973 by President Richard Nixon in response to rising concern over the number of species threatened by extinction, the ESA protects over 1,650 U.S. species by prohibiting negative impacts on species and their habitats and guiding the recovery of populations (2). Today, the ESA remains the primary piece of environmental legislation for protecting imperiled species and recovering them to the point that the law’s protections are no longer needed. With such a crucial role, the ESA must be implemented correctly. Yet agencies often struggle with gaps in effective implementation as they face funding shortfalls and staff limitations alongside a rising number of listed species. Although the ESA is a strong law, effective implementation in the face of these challenges is key. Taking advantage of opportunities for improvement in efficiency and effectiveness is crucial if the ESA is to continue preventing extinction and recovering species.

Section 7 of the ESA directs federal agencies to use their authorities to conserve listed species and is a key aspect of the law’s strength. Under section 7(a)(2), federal agencies (“action agency”) are instructed to consult with the U.S. Fish and Wildlife Service (FWS) or the National Marine Fisheries Service (NMFS) if any action authorized, funded, or carried out may jeopardize listed endangered or threatened species or destroy or adversely modify species’ critical habitat (for definitions see Box 1, Glossary). If an action agency initially concludes that the action is not likely to adversely affect species or their critical habitat, the agency must request Service concurrence on its finding. If the Service concurs, the consultation is complete; this assessment is classified as an “informal consultation.” Conversely, if an action is deemed likely to adversely affect species or critical habitat, a “formal consultation” is initiated, and the consulted Service will issue a biological opinion with their findings of the project’s impact on imperiled species. FWS and NMFS share administration of the ESA, with NMFS generally overseeing marine species and FWS managing terrestrial and freshwater species (3). However, both Services have authority over some listed species that cross jurisdictional boundaries, such as sea turtles, and consult with action agencies on these joint-jurisdiction species. If done properly, consultations ensure that federal agency actions do not violate the jeopardy and adverse modification prohibitions of the ESA, thereby minimizing negative effects on listed species.

### Box 1: Glossary

*Glossary of terms typically used to describe and discuss consultations under section 7(a)(2) of the U.S. Endangered Species Act. The exact legal and policy definitions can be found in the referenced Code of Federal Regulations (CFR) and Handbook sections.*

#### Action

All activities or programs of any kind authorized, funded, or carried out, in whole or in part, by Federal agencies in the United States or upon the high seas. [50*CFR§*402.02]

#### Action agency

The federal agency proposing the action.

#### Biological opinion

The document resulting from formal consultation that describes the proposed action, the Service evaluation of the effects of the action, the determination of whether the species’ existence is jeopardized or its critical habitat is adversely modified, and any conservation requirements for the action agency. [50*CFR§*402.02, 50*CFR§*402.14(*h*)]

#### Critical habitat

The specific areas and habitats essential to conserving the species. Critical habitat may be designated in areas that are occupied or unoccupied at the time of listing. Occupied habitat must also have “physical or biological features” that require special management considerations or protection. [*ESA§*3(5)(*A*)]

#### Formal consultation

The type of detailed evaluation undertaken for federal actions that are likely to adversely affect one or more ESA-listed species. [50*CFR§*402.02, 50*CFR§*402.14]

#### Informal consultation

The type of detailed evaluation undertaken for federal actions that are not likely to adversely affect one or more ESA-listed species. [50*CFR§*402.02, 50*CFR§*402.13]

#### Jeopardy (to jeopardize)

To engage in an action that reasonably would be expected, directly or indirectly, to reduce appreciably the likelihood of both the survival and recovery of a listed species in the wild by reducing the reproduction, numbers, or distribution of that species. [50*CFR§*402.02]

#### Programmatic consultation

A consultation that addresses multiple actions taken by an agency on a program, regional, or other basis. For example, programmatic consultations may cover many different energy development projects within particular Bureau of Land Management lands in a single, landscape-level evaluation. (Handbook, p. xvii)

#### Take

To harass, harm, pursue, hunt, shoot, wound, kill, trap, capture, or collect, or to attempt to engage in any such conduct [*ESA§*3(19)]

The consultation process is guided by the Section 7 Handbook (hereafter, Handbook), which was created by the Services to “promote efficiency and nationwide consistency [of consultations] within and between the Services” (4). The Handbook guides biologists to ensure consultations are serving their purpose of adequately protecting listed species to the fullest extent of the ESA and lays out a framework for what should be included in each section of a biological opinion issued by the Service. However, the Handbook is a guidance document only and does not prescribe all details of a consultation. This results in variation in consultation completeness, which could become problematic if differences introduce inefficiencies or inconsistencies that ultimately reduce the protection or conservation of imperiled species.

Two preliminary observations suggest consultation completeness may differ between the Services in ways that reduce consultation effectiveness. First, recent analysis of data on all section 7 consultations recorded by FWS from 2008-2015 (5) revealed discrepancies in the time duration of consultations between the Services. Whereas the FWS completed 80% of formal consultations within the 135-day time limit set by the Handbook (the proportion of on-time consultations is likely higher because the data do not include information on legitimate “pauses” during consultation; JWM and Y-WL, pers. obs.), NMFS completed only 30% in this timeframe (6). This discrepancy in timing could indicate a problem in the conservation process if, for instance, FWS is compromising quality of the analyses for quantity in order to complete its required number of consultations, which is substantially greater than NMFS despite receiving similar levels of funding (7; 8). Second, based on the authors’ combined experience of reading hundreds of consultation documents, we observed high variation in the general completeness and consistency of consultation documents (authors, pers. obs.). Variation appears to be structured (e.g., by species or office) rather than random, and especially large differences occur between consultations produced by the two Services. There are numerous reasons why the FWS and NMFS could differ in their approach to or process for consultations. For example, the two agencies have overlapping but not identical legal mandates; different organizational histories and cultures; and receive different levels of funding, differences that percolate across regions and offices within each Service (9). Understanding the type and degree of variation among consultations could help identify the cause and outcome of differences. That knowledge can in turn assist in designing solutions that minimize inconsistencies and maximize quality of the consultation process to support the Services in enforcing the ESA. Yet to our knowledge, there has never been a systematic analysis of differences in consultation completeness, creating a knowledge gap with direct implications for biodiversity conservation and environmental policy.

Here we quantify and evaluate variation in how the Services implement section 7 by comparing the completeness of consultation documents for threatened and endangered species of sea turtles against the requirements of the Handbook. Sea turtles are one of the few taxa which falls under the jurisdiction of both the FWS and NMFS, offering a unique opportunity for direct comparison of consultation completeness. As we discuss further below, we expect consultations that follow the requirements of the Handbook are more complete and more likely to result in better conservation outcomes because the Handbook provides the best available description of how to comply with section 7. Thus, we assess completeness of a consultation under the assumption that a more complete document will lead to better conservation for the species. In doing so, we take advantage of a natural experiment to analyze the differences in how the Services implement the consultation process. While the null hypothesis may be equality of consultation document completeness, based on previous observations, we expect NMFS consultations to more complete than FWS consultations. We report significant differences in the completeness of both the formal and informal consultations between the Services. Our results highlight several pathways by which the Services can systematically improve the completeness and quality of consultations to strengthen the ESA and improve the protection and recovery of North America’s most imperiled species.

## 2. METHODS

### 2.1 Sampling

The Services have carried out hundreds of thousands of consultations since the ESA was established. Because consultations are often context-specific and can differ depending on specific categories such as action type and species, fully random sampling of species was not suitable for our objective. Following prior methods (10), we chose a defined subset of consultations to make comparisons between the Services more direct and insightful. We controlled for extraneous sources of variation by conducting our analysis on consultations from January 2008 through April 2015 and involving actions proposed by the Army Corps of Engineers (the Corps) that could potentially impact sea turtles in Florida. This focus enabled us to minimize confounding factors that might be introduced by the time period, type of action being evaluated, or species natural history or geographic variation, and therefore to focus on differences between the Services’ consultation process and output. Species of sea turtle were the most consulted on by the Corps and included green sea turtle [*Chelonia mydas*], loggerhead sea turtle [*Caretta caretta*], Kemp’s ridley sea turtle [*Lepidochelys kempii*], leatherback sea turtle [*Dermochelys coriacea*], and hawksbill sea turtle [*Eretmochelys imbricata*].

### 2.2 Consultation Selection

We obtained consultation data that met our sample criteria from several publicly available databases. We accessed NMFS consultations using the Public Consultation Tracking System (PCTS; https://pcts.nmfs.noaa.gov/pcts-web/homepage.pcts), which allows users to directly download consultations. FWS has a similar database of consultation records, the Tracking And Integrated Logging System (TAILS). TAILS is designed to help coordinate record-keeping between field and regional offices of FWS and does not provide the consultation documents. Instead, the TAILS database provides records of FWS consultations but has no public interface, therefore we accessed TAILS records using the Section 7 Explorer web application (https://defenders-cci.org/app/section7_explorer; Malcom and Li 2015) that allows the public to search for consultations using TAILS data. Using PCTS and the Section 7 Explorer to identify the set of consultations involving the Corps and sea turtles, from which we randomly selected 30 formal and 30 informal consultation records from each Service during the study time period. We acquired the NMFS consultations directly from PCTS, while those from FWS we acquired through FWS South Florida Field Office’s online document library for biological opinions (https://www.fws.gov/verobeach/verobeach_old-dontdelete/sBiologicalOpinion/index.cfm) or through a Freedom of Information Act (FOIA) request. While evaluating the original selection of NMFS formal consultations, we discovered some that did not assess sea turtles in the biological opinion despite search parameters constrained to sea turtles. To account for this discrepancy, we removed those not assessing sea turtles and randomly selected an additional 10 formal NMFS consultations for evaluation from the PCTS database. All of the consultations analyzed in this work are archived at Open Science Framework (OSF) under https://dx.doi.org/10.17605/OSF.IO/KAJUQ.

### 2.3 Evaluation Criteria

We recorded the start and end dates of the consultation, year completed, regional office filed through, species of sea turtles, page length, and other general information for each consultation. All evaluated consultations and data are provided at OSF (https://dx.doi.org/10.17605/OSF.IO/KAJUQ). We developed different scoring methodologies for formal and informal consultations because each type involves different content as detailed in the Handbook. Scoring rubrics are provided in SI Appendix 1 (formal consultations) and Appendix 2 (informal consultations). It was not feasible to blind scorers to the Service that wrote consultations because of the nature of the documents; any familiarity with the consultation process makes the Service immediately apparent. Therefore, reviewers were not blind to the Service when analyzing completeness. When there was any ambiguity as to the appropriate score, a second reviewer (JWM) would read the consultation in question, then decide on the appropriate score with the primary reviewer (ME).

For formal consultations, we selected the four core sections from the Handbook to score the completeness of each biological opinion: “Status of the Species,” “Environmental Baseline,” “Effects of the Action,” and “Cumulative Effects.” Although not an exhaustive list of biological opinion sections, these four sections contain the bulk of the information and analysis of the species and proposed action. The Status of the Species and Environmental Baseline sections received a score from 0-5 and the Effects of the Action and Cumulative Effects sections were given a score from 0-2 based on how well they met the specific requirements for that section by the Handbook. Rating the completeness of these core sections of the biological opinion was straightforward because the criteria described by the Handbook allowed for a simple present/absent scoring system. For some analyses, these present/absent scores were summed for each of the four core sections. We also calculated total completeness by summing the scores across all four sections. The overall completeness was normalized by calculating the ratio of the summed score to the total points possible for each consultation.

Scoring the informal consultations used a simpler rubric because informal consultation documents are shorter, rarely have individual sections, and the Services generally do not prescribe the required contents. We surveyed a selection of informal consultation documents from both Services and considered what information Services personnel need in order to evaluate the effects of actions and monitor the action after consultation is complete. We identified five criteria to evaluate the completeness of informal consultations: stating the action, analysis of the action, analysis of the impacted species, stating the reason why the consultation stayed informal and including a map of the area affected by the action. Though a map is not required by the Handbook, the action area is highly important for much of the consultation analysis, and thus the inclusion or omission of a map was scored. These criteria were each assigned 1 point, for a total possible score of 5 points.

During preliminary work on this project we noticed the use of “sticker concurrences,” in which the FWS South Florida Office recorded only a sticker of consent applied to the request for concurrence provided to FWS (SI Figure 1). This sticker of approval for the action was in lieu of a complete informal consultation, and no additional consultation documentation was supplied. Despite their lack of analysis, sticker concurrences were scored in the same manner as all other informal consultations.

### 2.4 Statistical Analyses

Our goal was to understand patterns and associations of variation in consultation completeness. We used summary statistics (mean and standard deviation) and Pearson’s correlations to describe patterns. To examine relationships between completeness and associated factors, we used two modeling approaches: a binomial generalized linear model (GLM; 11) to identify predictors of the proportions of total possible points, and ordinal logistic regression (OLR; 12) to analyze the individual component scores. We considered six variables that were most likely to affect consultation completeness: the Service performing the consultation, whether the consultation was formal or informal, the year the consultation took place, the species of sea turtle assessed, the type of action assessed, and whether the consultation was part of a programmatic consultation (see Glossary). We incorporated these variables into a global model (Model 1) of all variables and eight additional subset candidate models for the analysis of overall completeness using the GLM (Table 1). We also considered that the particular office within the Service might be an important predictor of consultation completeness. However, given that our focus is on the potential differences between the Services and that the offices are nested within the Services, the office variable was not included in our candidate model set. Because of the fundamental differences between formal and informal consultations and the difference in total possible score, we calculated the response variable as the proportion of possible points for each consultation. When we analyzed data separately for formal and informal consultations, we used reduced candidate model sets by removing the informal consultation variable from formal analyses and the formal and programmatic variables from the informal analyses.

**Table 1.**
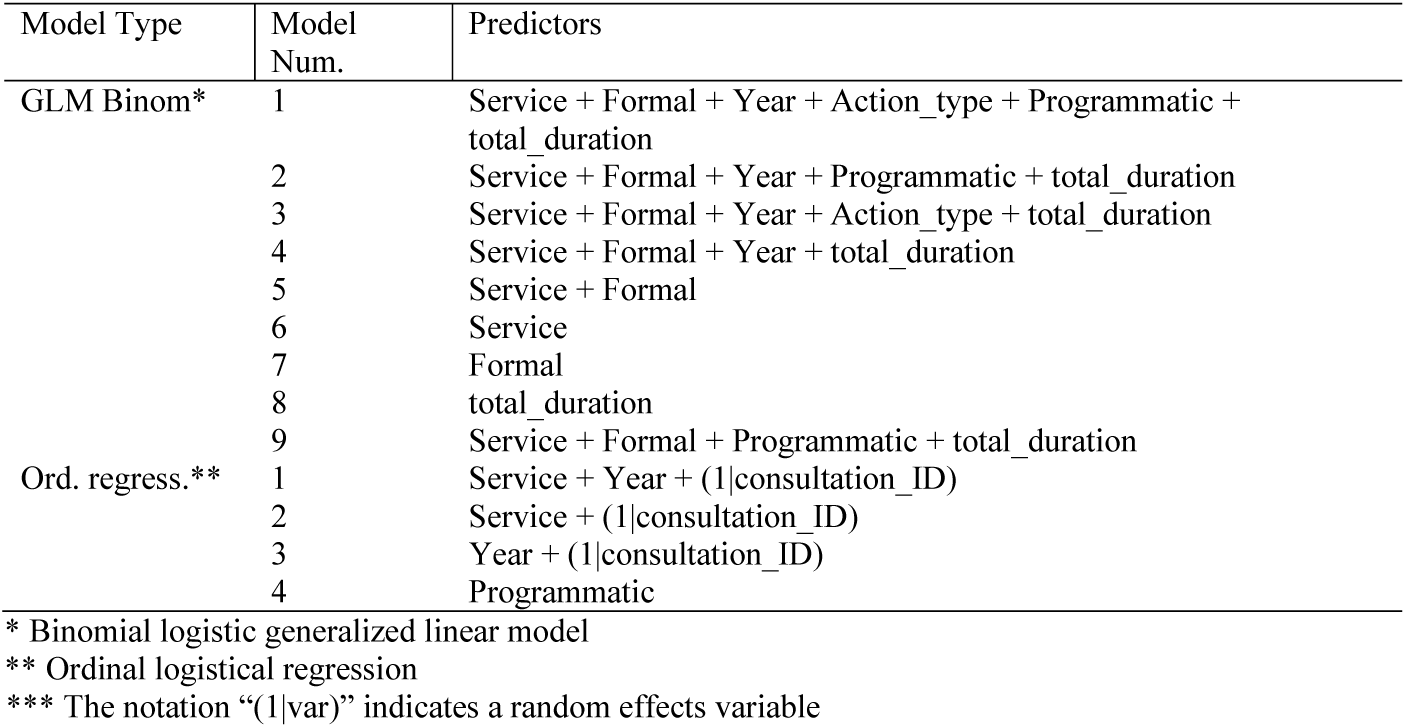
Candidate generalized linear and ordinal regression models for predicting overall consultation completeness and conservation action specificity.

We used a set of three candidate ordinal regression models (Table 1) with random effects for the consultation document in which the components were nested. While programmatic consultation was an important predictor of completeness in the overall analysis, the Hessian was singular (presumably because of the lack of NMFS programmatic consultations) for the components and we were not able to include programmatic as a variable in these analyses. We therefore evaluated summary statistics to investigate the role of programmatic consultations in shifting completeness scores. We used the R package ‘ordinal’ (13) to conduct ordinal regression. A univariate analysis was performed to identify predictor variables.

We carried out model selection (14) based on Akaike’s Information Criterion adjusted for small sample sizes (AIC_C_) using the AICcmodavg package (15). We considered models with ΔAIC_c_ > 2.0 as having strong support (14). All analyses were done in R 3.3 (16) and are available as a package vignette in the project’s OSF repository (https://dx.doi.org/10.17605/OSF.IO/KAJUQ).

### 2.5 Consultation Process

To supplement data gathered from the consultation documents, one of the authors (ME) discussed the consultation process with one biologist from NMFS and six biologists from FWS who consulted on sea turtles in Florida. These biologists were on the list of Service personnel who worked directly on the consultations evaluated for this study and were selected based on availability. Information collected on the consultation process was not meant to be representative of a larger sample but was instead intended to provide further insight into results. Biologists were asked about the consultation process concurrent with our scoring of the consultations (in August 2015) at the agency offices in Florida. The questions were based on our understanding of the Handbook and preliminary examination of the consultations we reviewed. We asked biologists about their opinions on the consultation process and how well consultations serve the intended purpose (SI Appendix 3). We then coded answers into categories of similar themes. All biologists were spoken to under the condition of anonymity and with full awareness of the agencies. Informed consent was obtained from all participants. Although the sample size is too small for statistical analysis, we reviewed and scored the notes on the consultation process from the biologists to summarize recurring themes.

## 3. RESULTS

We retrieved, read, and scored 55 consultations produced by FWS (30 formal and 25 informal) and 68 consultations produced by NMFS (38 formal and 30 informal) for a total of 123 consultations. Consultations assessed the effects of the action on seven species on average (Table 2). Formal consultations ranged in length from 1 to 120 pages and required over a year to complete on average. Of the core completeness sections evaluated, ‘Status of the Species’ was by far the longest, with an average of 19 pages. This section often contained lengthy content that was neither relevant to the species’ life history in the geographic area of the action nor to the effects of the action. In our random sample of FWS informal consultations, only one featured the sticker concurrence that we observed in the preliminary work.

**Table 2.** Summary statistics across all 123 formal and informal consultations.

| Consultation type | Variable | Mean | Min | Max | SD | N* |
| --- | --- | --- | --- | --- | --- | --- |
| Formal | Length (pages) | 34.6 | 1 | 120 | 21.1 | 284 |
|  | Duration (days) | 371.5 | 6 | 1691 | 320.2 | 340 |
|  | No. of species (total) | 7 | 4 | 18 | 3.6 | 324 |
|  | No. of References | 164.3 | 1 | 434 | 121.4 | 330 |
|  | Species Status length (pages) | 18.7 | 0 | 67 | 12.5 | 325 |
|  | Baseline length (pages) | 6.7 | 0 | 23 | 4.7 | 318 |
|  | Effects length (pages) | 5.4 | 0 | 15.5 | 3.9 | 303 |
|  | Cumulative Effects length (pages) | 0.7 | 0 | 1.5 | 0.3 | 298 |
|  | CR** | 0.9 | 0 | 1 | 0.3 | 292 |
|  | CM** | 0.5 | 0 | 1 | 0.5 | 272 |
|  | RPM** | 0.8 | 0 | 1 | 0.4 | 287 |
| Informal | Duration (days) | 163 | 0 | 1227 | 223.3 | 260 |
|  | No. of species | 7.0 | 1 | 49 | 6.0 | 265 |
|  | Construction Conditions | 0.7 | 0 | 1 | 0.4 | 264 |

\* Numbers are based on individual turtle species per consultation because the jeopardy and adverse modification conclusion is made on per-species basis for an action. \*\* CR = Conservation Recommendations made by the Services; CM = Conservation Measures proposed by the action agency; RPM = Reasonable and Prudent Measures to minimize the amount of take resulting from an action.

### 3.1 Overall Consultation Completeness

Generalized linear modeling suggested that consultation completeness was best explained by Model 9, which showed the lowest AIC_c_ (ΔAIC_c_ = ∼2; Table 3). This model, which included all predictors except action type and year, indicated that a consultation done by NMFS was 1.40 times (95% CI = 1.25 - 1.57; Figure 1a) as likely to receive a higher score for completeness as a consultation done by FWS. FWS’s programmatic consultations provided a significant completeness boost (OR = 1.35; 95% CI = 1.17 - 1.56), but formal consultations were about as likely (OR = 1.0; 95% CI = 0.89 – 1.13; Figure 1b) to score higher as informal consultations (Table 4). The duration of consultations was positively associated with overall completeness in a univariate GLM (*r* = 0.20; *p* = 1.04e^-6^) but did not rank as an important variable in the multivariate analysis. Similarly, the section length in pages was also correlated with completeness in a univariate analysis (r = 0.2, *p* = 0.0037). However, after accounting for the Service performing the consultation and for programmatic consultations in a binomial GLM, there was no relationship (*z* = 1.024, *p* = 0.306). Model 2, which included the same predictors as Model 9 but added in the year the consultation was completed, was also supported. This model indicated that the year was associated with a slight decrease in consultation completeness over the study period, though this association was not statistically significant (OR = 0.993; 95% CI = 0.97 – 1.02), thus we focus on model 9.

**Fig 1.**
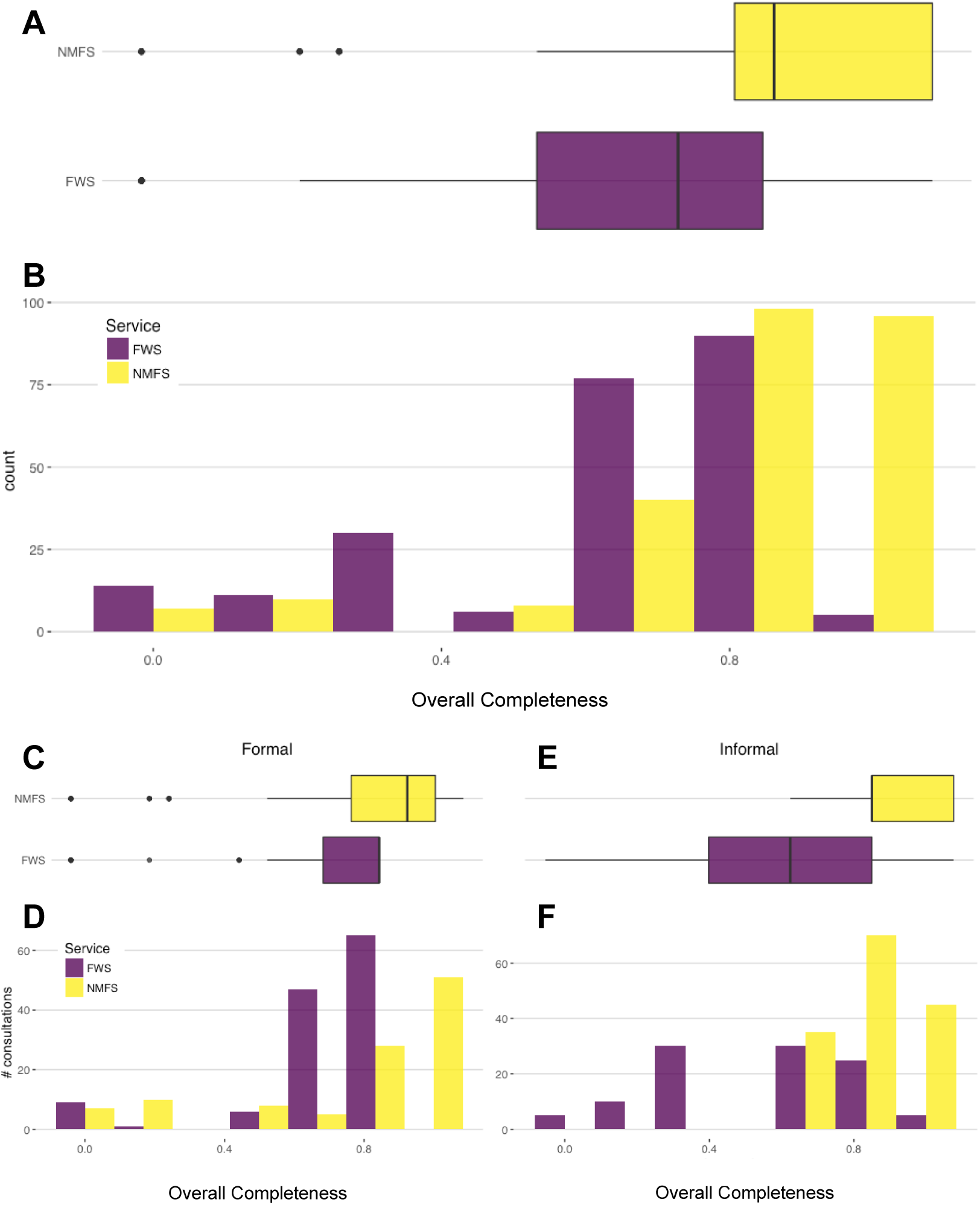
Completeness scores for NMFS consultations were higher on average than scores for FWS consultations across all consultations (A), formal consultations (B), and informal consultations (C). The overall completeness score for each consultation is the sum of points scored divided by the sum of points possible (see Methods for details). *Top panel:* Histogram and boxplots of all consultations (formal and informal, including programmatic consultations) for each Service. *Bottom panel:* Overall scores plotted by Service for formal and informal consultations separately.

**Table 3.**
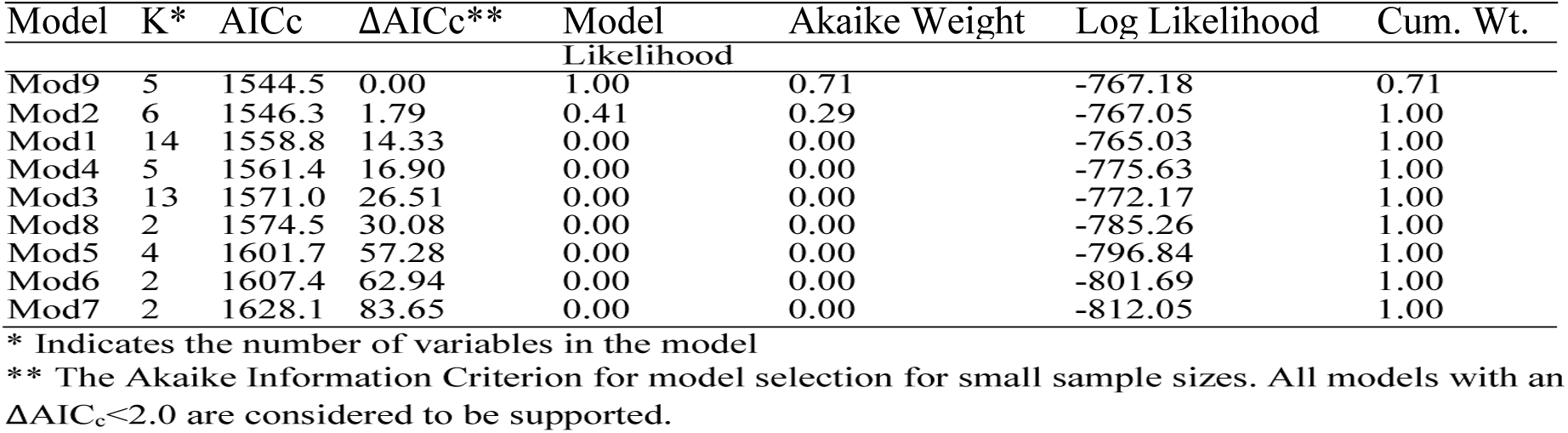
Generalized linear model selection results for overall completeness across 123 FWS and NMFS consultations.

**Table 4.** Odds ratios (OR), confidence intervals, and parameter statistics for model 9, the best-supported candidate set for predicting overall consultation completeness.

|  | OR | LCL (2.5%)* | UCL (97.5%)** | Model z-value | p-value |
| --- | --- | --- | --- | --- | --- |
| (Intercept) | 5.54E<sup>-01</sup> | 4.93E<sup>-01</sup> | 6.23E<sup>-01</sup> | -9.883 | 4.94E<sup>-23</sup> |
| Service (NMFS) | 1.40 | 1.25 | 1.57 | 5.689 | 1.28E<sup>-08</sup> |
| Formal (yes) | 1.00 | 0.89 | 1.13 | 0.042 | 9.66E<sup>-01</sup> |
| Programmatic (yes) | 1.36 | 1.18 | 1.57 | 4.202 | 2.64E<sup>-05</sup> |
| total duration | 1.00 | 1.00 | 1.00 | 1.454 | 1.46E<sup>-01</sup> |

\* LCL = Lower control limit
\*\* UCL = Upper control limit

### 3.2 Components of Completeness

We examined sources of variation in the components of overall consultation completeness. The only component of formal consultations that exhibited a strong association with any predictor variables was the Environmental Baseline, for which Service was a strong predictor of completeness and NFMS was more likely to produce more complete consultations (*z* = 5.3993, *p* = 6.691e^-8^; OR_NMFS_ = 2.6e^4^ [95% CI = 6.5e^2^ – 1.1e^6^]; Figure 2). For the Environmental Baseline section, NMFS consultations were more comprehensive and tended to include previous consultations in the action area and discuss critical habitat or lack thereof as per the Handbook. Neither of these characteristics were consistently present in FWS consultations. Most of the completeness components of informal consultations were similar except for two categories (Figure 3). The analysis of the action and the reason the consultation was informal were associated with the time duration of the consultation (at a nominal α = 0.05): generally, the longer the informal consultation took to complete, the more likely these components were included. Second, although not required by the Consultation Handbook, half of NMFS but only 15% of FWS informal consultations included a map of the proposed action.

**Fig 2.**
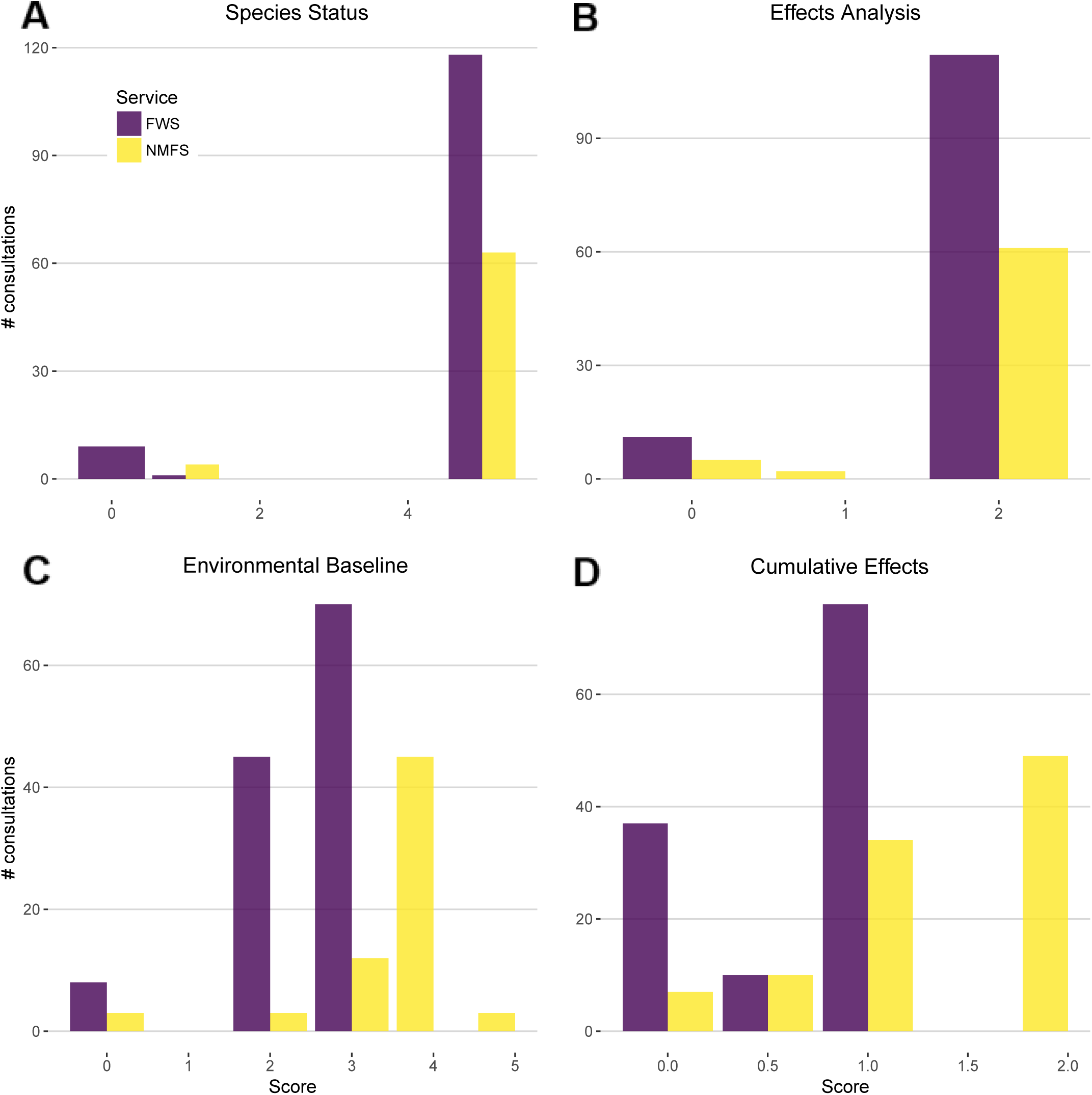
Individual components of consultations produced by NMFS showed higher completeness scores than those by FWS on average. However, the only component that statistically differed between the Services was the Environmental Baseline (*z* = 5.3993, *p* = 6.691e-08; OR_NMFS_ = 2.6e^4^ [95% CI = 6.5e^2^ – 1.1e^6^]). The scores are the raw completeness scores for formal consultation components.

**Fig 3.**
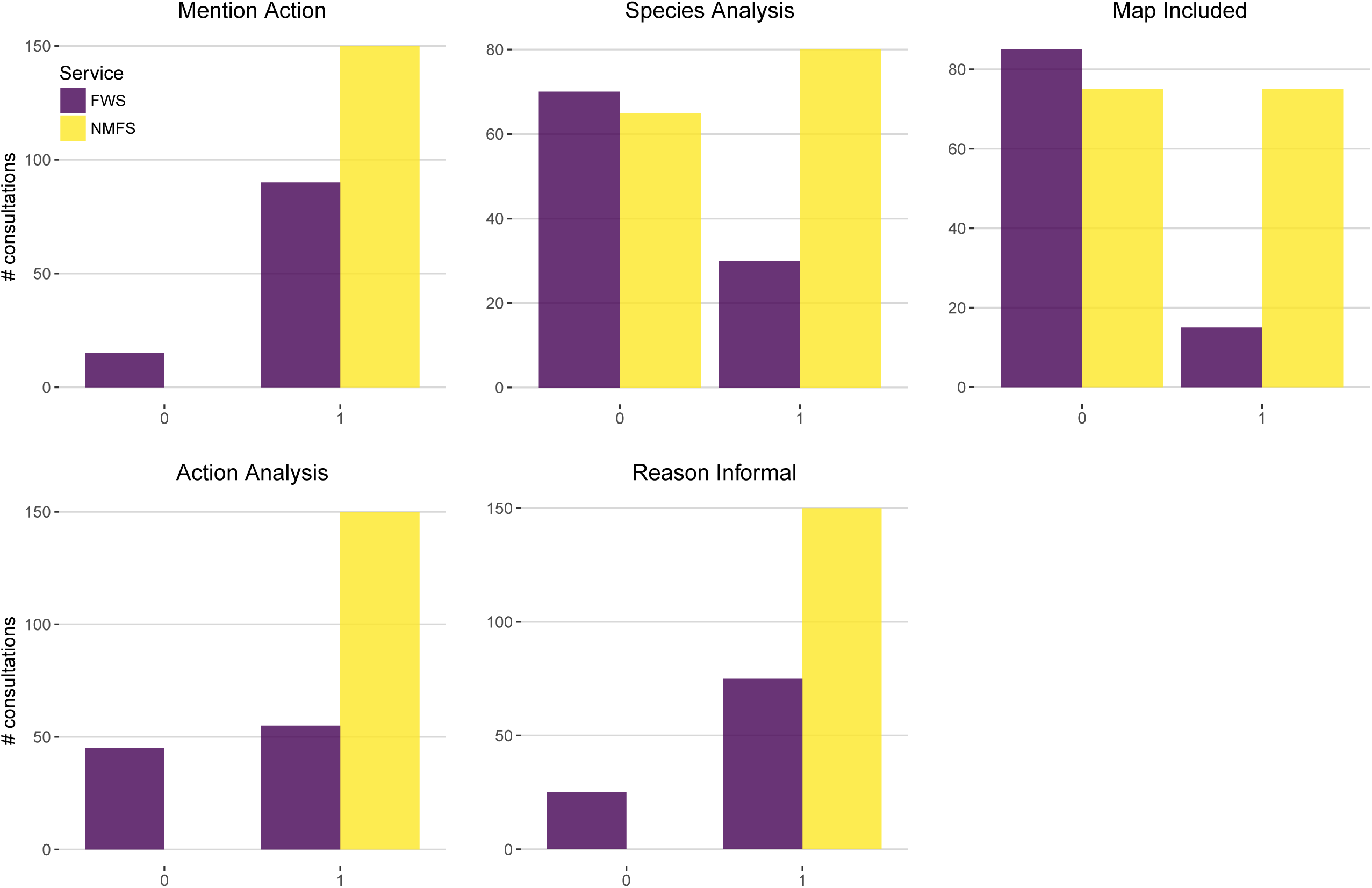
Informal consultations from NMFS featured more information and therefore showed higher completeness scores than those from FWS on average. The components of informal consultation completeness scores were binary (0 indicates absence; 1 indicates presence) in the consultations.

### 3.3 Consultation Process Feedback

We spoke with six biologists from FWS and one from NMFS and coded their responses into categories of similar themes (Table 5; full response notes in SI Appendix 4). When asked how the consultation process could be improved, most biologists (6 of 7) mentioned they found the process frustrating and many stated that they were overwhelmed with work. One biologist pointed to the fear of possible litigation resulting from shorter consultations as a reason for the overly comprehensive and highly time-consuming consultations that are currently the norm. Five of seven biologists also favored expanding the use of consultation keys, which are designed to help the biologists improve the timing and consistency of consultations when appropriate for a species or on a case-by-case basis (see, e.g., http://www.fws.gov/panamacity/resources/WoodStorkConsultationKey.pdf; SI Appendix 5). All biologists except one mentioned that they keep a record of cumulative incidental take, which varied in form from notes kept on a whiteboard to Excel spreadsheets. However, only three consultations (all from NMFS) incorporated a tally of previously authorized take in the analysis of the effects of the current action on sea turtle populations.

**Table 5.** Responses to a selected sample of consultation process questions asked of FWS/NMFS biologists.

| Biologist | Favor consultation keys | Often encounter scientific uncertainty | Tally cumulative take | Frequently reference section 7 Handbook | Favor publicly available consultations | Suggestions for improvement |
| --- | --- | --- | --- | --- | --- | --- |
| 1 | In some cases | No | Yes | Yes | Yes | Inter-office consistency |
| 2 | Yes | No | Yes | No | Yes | None |
| 3 | No | No | Yes | Variable | Yes | Inter-office consistency |
| 4 | Yes | Rarely, assume species is present | Yes | No | Yes | Intra- and inter-office consistency |
| 5 | In some cases | Rarely, assume species is present | Makes an attempt | Yes | Yes | BiOp streamlining |
| 6 | In some cases | No | Yes | Yes | Yes | Inter-office consistency |
| 7 | No, too nuanced | Yes, defer to species | No - too difficult | No | Yes | Improve efficiency |

## 4. DISCUSSION

The ESA is considered one of the strongest wildlife protection laws in the world (17), and section 7 is a foundation of this strength. The content and quality of section 7 consultations can alter conservation outcomes, but such protections can only be realized if the scientific and regulatory analyses are robust. Despite the importance of consistently high-quality consultations, no analyses have critically evaluated the strengths and weaknesses of these regulatory documents. Our analysis offers an urgently needed first step towards understanding the quality of consultations to inform and improve future consultations. Across all 123 consultations evaluated, we found that completeness relative to the standards in the Handbook varied significantly between the Services: NMFS consultation documents were consistently more complete than FWS consultation documents. We interpret this difference in content as a difference in consultation quality that may be affecting the conservation of ESA-listed species. In combination with the biologist discussions, which illuminate some of the possible causes of variation, our results reveal specific areas of improvement to ensure that future consultations achieve their objective of protecting threatened and endangered species.

### 4.1 Consultation Quality

The completion of both formal and informal consultations was higher in documents produced by NMFS than FWS. This result is consistent with prior findings that NMFS scored higher than FWS in three of seven metrics characterizing the use of “Best Available Science” in recovery plans, lawsuits, listing decisions, and literature cited in biological opinions and no difference was detected between the agencies in the other four metrics (9). Although the cause of the difference is beyond the scope of our study, our discussions with Service biologists suggested one possible explanation: that the lack of time and resources available for the agencies’ ever-increasing consultation workload may limit their quality. The FWS biologists especially stressed this point, which reflects the funding shortfall experienced by the FWS endangered species program. This program receives approximately equal funding as the Office of Protected Resources at NMFS even though Ecological Services within FWS is responsible for 15 times as many ESA-listed species (9). Expenditures per consultation is therefore likely much lower for FWS. Future research should investigate how the Services allocate funding to consultations compared to other endangered species program components, such as listing and recovery.

Our scoring of the individual sections of biological opinions provides further insight into why FWS consultations are lower completeness than NMFS consultations and for which content both Services deviate from the expectations of the Handbook. Although documents by both Services consistently showed low completeness in the Environmental Baseline section because previously authorized incidental take in the action area was rarely analyzed, FWS scored lower than NMFS because the take analysis was missing from all prior consultations. The lack of this analysis is one of the most pernicious problems with implementing the ESA (10). The omission of hundreds or thousands of minor take actions from analysis in consultations can compound to result in “death by a thousand cuts,” whereby individual actions are insignificant for the species but the cumulative effects across many actions severely damage their populations (18). A 2009 Government Accountability Office report on FWS’s implementation of the ESA highlighted this concern and recommended that the Services track authorized take across a species’ entire range to better inform consultations (19). The only three consultations that included an analysis of previously authorized take were all produced by NMFS, enhancing the difference in completeness between the Services for this core section. However, it is worth noting that FWS’s programmatic consultation for beach work across Florida (Activity Code 41910-2010-F-284) listed previous formal consultations. Unfortunately, those data were not analyzed in the evaluated consultation and there was no evidence they played a role in the Environmental Baseline or the Effects Analysis. It is unclear why previously authorized take in the action area was not analyzed, especially since many biologists that we spoke with stated that they record cumulative take. Future research should investigate the disconnect between the information that Services biologists record and the information included in consultations.

Although the Handbook requires certain analyses for each section, sections of many FWS consultations contained little or no analysis and instead merely repeated the boilerplate language from the Handbook. This was particularly true of the Cumulative Effects section of FWS consultations, which often mentioned the obligation to “include the effects of future State, tribal, local or private actions that are reasonably certain to occur,” followed by a statement that there would be no cumulative effects. In contrast, most NMFS consultations more thoroughly analyzed the cumulative effects, which are critical to understanding the effects on species conservation status.

The Handbook guidance for informal consultations is less prescriptive than for formal consultations, but our analysis revealed that the completeness of consultations by FWS is similarly lower than for NMFS. Three components — the analysis of the action, the species analysis, and a map of the action area — were consistently missing or insufficient in the informal FWS consultations that we reviewed. On one hand, because informal consultation is merely a prerequisite to determine whether formal consultation is warranted, we recognize that detailed informal consultation analysis is unlikely to benefit ESA-listed species. Nonetheless, omission of content means that the administrative record is inconsistent and incomplete (see ref. 20 for a relevant discussion) and, most alarming of all, differs from the Services’ expert recommendations for informal consultations. This is apparent in the use of “sticker” concurrences, observed both in our preliminary work and in one randomly sampled informal consultation. While these stickers may save time, they provide no record of why FWS approved the action or method for assessing whether FWS properly implemented that component of the ESA. Furthermore, in contrast, all informal consultations from NMFS explained why the consultation was informal. The shortcomings of FWS informal consultations can likely be explained by the resource constraints, yet we highlight this example as an invitation for the agency to critically evaluate whether such shortcuts appropriately achieve greater efficiency, or whether different improvements could make the process more effective.

### 4.2 Opportunities for Improving Consultation Efficiency

The stark difference between the FWS and NMFS in consultation completeness highlight gap in the way section 7 is implemented. This discrepancy, coupled with the known disparity in both workload and resources (both financial and personnel) available per consultation, means that improving the efficiency with which the Services carry out consultations is essential to properly implementing the ESA. Ideally, the Services should spend enough time on each consultation so as to maximize the conservation benefit to a listed species. Awareness of this optimal threshold, and the required content to reach it, would avoid overspending precious resources (21). Here we discuss some critical inefficiencies, and potential pitfalls of efficient approaches, indicated by our results.

The higher completeness scores associated with consultations tiered off of the FWS programmatic consultation indicate that programmatic consultations are one promising way to improve consultation efficiency. The effects analysis of programmatic consultations should provide a better description of cumulative effects because many planned or potential projects within a program are evaluated together rather than individually. We expect that when the cumulative impacts are properly acknowledged, the assessment of jeopardy or adverse modification is more likely to reflect real-world conditions. Another benefit is that because the overall program has already been evaluated, the consultations for future individual projects are faster and can contain less analysis. Malcom and Li (2015) found that project-level consultations that tiered off of a program-level consultation were completed nearly three times faster than the average standard consultation. In the set of consultations we evaluated, the single FWS program-level programmatic consultation for beach renourishment across Florida was a “tide that raised all boats,” in which the project-level programmatic consultations that tiered off of the program-level programmatic consultation “inherited” the (generally) high scores of the program-level consultation and significantly increased the completeness of FWS consultations. Whether this is an outlier or representative of programmatic consultations in general is unclear but deserves further investigation. But the converse is also possible: low-quality program-level programmatic consultations would mean that tiered consultations inherit low-quality analyses that would likely lead to poor conservation outcomes. While the results from this set of consultations are promising, the Services need to continually evaluate their programmatic consultations to ensure that the speed benefits of these consultations do not overshadow the need for high-quality analyses.

Our discussions with biologists from the Services provided important context for interpreting the results and indicated other possibilities for improving consultation efficiency. The lack of consistency among offices and between Services was frequently mentioned as a frustrating aspect of the consultation process. The differing approaches to consultations can be difficult for action agencies as well, who can see the approval of a project depend largely on the consulting office (Y-WL and JWM, pers. obs.). One possible solution that we did not test is the use of consultation keys, as have been developed for Army Corps of Engineers consultations for a few species, including wood storks (*Mycteria americana*) and indigo snakes (*Drymarchon couperi*). The Services use these documents to promote appropriate standards for certain construction activities. Creating similar documents for other frequently-consulted species may streamline consultations and increase inter-office and inter-Service consistency. The use of consultation keys would also increase the transparency of the consultation process, making it easier for action agencies or their applicants to plan their projects.

We note one particular aspect of consultations that was not amenable to quantitative analysis but suggests efficiency improvements: inclusion of extensive material seemingly irrelevant to evaluating the effects of the action. For example, several consultations we reviewed included >20 pages of information on red knots (*Calidris canutus*), of which one paragraph was relevant to evaluating the action (JWM, pers. obs.). Including such inconsequential background information requires additional time not only for Services’ biologists, but also for the action agency or their applicants who read the opinion. By way of explanation, one FWS biologist mentioned that such information was included to buffer against any potential legal action, ensuring all “bases are covered.” However, this approach conflates “more” with “better” — the added time and cost does not always produce commensurate benefits for legal defensibility or conservation (22). We encourage the Services to critically evaluate the information in biological opinions and exclude irrelevant material. The Recovery Planning Initiative (RPI) now being adopted by FWS (SI Appendix 6) can help with this extraneous information problem. One component of RPI is a single, continually updated Species Status Assessment (SSA) for each ESA-listed species, which would be incorporated by reference in consultations, conservation permits, five-year reviews, and other aspects of ESA implementation (SI Appendix 7). Widespread adoption of SSAs would improve efficiency and, because they should include an analysis of previously authorized take, improve the effectiveness of section 7 consultations.

A simplifying assumption we made is that a more complete consultation that addresses each of the parameters of the Handbook will lead to better conservation outcomes for the species and is thus a higher quality document. While not every parameter set by the Handbook will help advance the goal of the consultation equally, addressing each parameter is important for understanding the rationale of the Service and action agency throughout the evaluation process. For these reasons, we believe the completeness of the consultation document holds substantial importance for species conservation. A caveat to this methodology is that in reducing complex documents like biological opinions to a few indicators often means some nuances to individual situations are lost. This is inevitable in the translating of a qualitative document to a quantitative process, but in equally applying guidance from the Handbook, we avoid this to the best of our abilities.

### 4.3 Policy Recommendations

Our results provide a basis for several policy recommendations that would improve the Services implementation of section 7 of the ESA:

1. *Develop and require the use of a single database for recording and querying authorized take.* The component most commonly missing from consultations we reviewed was an analysis of previously authorized take in the action area. This is not surprising because FWS and NMFS have not yet established a unified, systematic way for their biologists to record authorized take, much less to comprehensively quantify and track previously authorized take to use in the jeopardy and adverse modification analyses. A centralized take database was recommended by the GAO over a decade ago (19) but has not yet been implemented by the Services. Implementing this recommendation would dramatically improve the completeness of the Environmental Baseline analysis of consultations. In turn, we expect better conservation outcomes for consulted-on species. In addition to consultations, an authorized take database would be invaluable for informing ESA-required five-year status reviews, such that harmful effects from consultations can be compared to beneficial effects from conservation activities.
2. *Establish a systematic review protocol to ensure that programmatic consultations, which can increase efficiency, do not reduce the effectiveness of consultation.* Programmatic consultations can increase consultation effectiveness and efficiency – in theory – but the Services must ensure that the quality of project-level consultations is not sacrificed. In our results, the programmatic consultation was the “rising tide that lifted all boats.” Ensuring that other and future programmatic consultations are similarly well-crafted can result in high quality, consistently-implemented consultations. The Services have expressed an interest in increasing the use of programmatic consultations and recently promulgated new regulations to do so (50 CFR § 402.14), but such an increase must formally guard against a loss of effectiveness. Regular reviews at the field office, regional, and national levels, guided by a robust “checklist” of effectiveness measures, could also benefit an expansion of the use of programmatic consultations.
3. *Require more widespread development and use of consultation keys.* Our results revealed variation in consultation completeness between the Services. If we had chosen a wider selection of consultations, this variation may have further increased. This highlights the need to promote standardization as a means of improving the efficiency and effectiveness of consultations. The biologists we spoke with suggested that the use of consultation keys could improve consistency. Although not every species and every type of action is amenable to consultation keys, wider use of keys could significantly improve the parts of consultations where they are relevant.
4. *Reduce workload by referencing prior documents.* To reduce the rote workload for consultation biologists and consulting agencies, the Services could consider transitioning to referencing SSAs, created as part of the Recovery Planning and Implementation strategy, in consultations. This would dovetail with FWS’s current revision of the recovery planning program, which places SSAs as a central piece of the process. Improving efficiency through standardization should not mean cutting corners, however. The informal concurrence stickers are a form of standardization, but, as currently used, they do not provide an adequate record of why decisions were made. They may be sufficient if modified slightly, such as by adding simple check boxes and short note fields to indicate the reason a consultation qualified as informal.

Implementing the above recommendations could significantly increase efficiency to better use the precious resources of the Services, and thus would improve the conservation benefit conferred by section 7 consultations. Strengthening the completeness of the consultations through these methods would enable the Services to improve the overall effectiveness of the ESA, thereby reinforcing its critical role in conserving imperiled species.

## ACKNOWLEDGMENTS

We thank the personnel from the Florida offices of the U.S. Fish and Wildlife Service; the St. Petersburg office of the National Marine Fisheries Service; and the Florida Fish and Wildlife Conservation Commission for their work on consultations and for the insights they provided us during this project. This research did not receive any specific grant from funding agencies in the public, commercial, or not-for-profit sectors.

---

**SI FIGURE 1.**
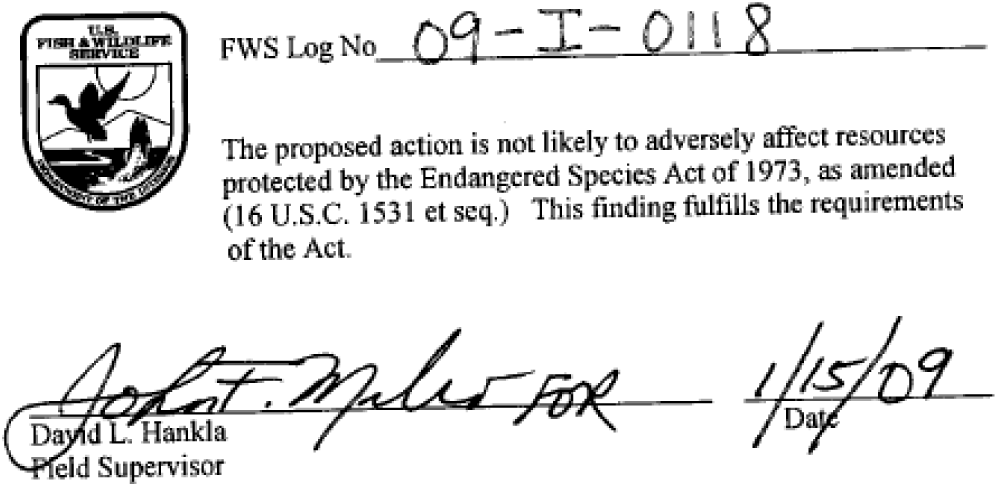
INFORMAL STICKER CONCURRENCE.

Complete informal consultation included in Open Science Framework archive at https://dx.doi.org/10.17605/OSF.IO/KAJUQ. Note that there is no accompanying analysis to clarify why this informal consultation was found not likely to adversely affect the species or any listed critical habitat.

## SI APPENDIX 1: SCORING RUBRIC FOR FORMAL ESA SECTION 7 CONSULTATIONS

### Environmental Baseline Completeness (Total Points: 5)

1. Does the Environmental Baseline address the status of the species in the action area? (1)
2. Is there a mention of past/ongoing threats to the species in the action area? (1)
3. Does the Environmental Baseline take past consultations in the action area into consideration? (1)
4. Is there mention of critical habitat (or lack thereof) for the species? Does said critical habitat overlap with the action area? (1)
5. Does the baseline include State, tribal, local and private actions already affecting the species that will occur contemporaneously with the consultation in progress, as per the handbook? (1)

### Effects of the Action Completeness (Total Points: 2)

1. There is a clear and defined cause and effect analysis of the action. (1)
2. The consultation gives an explanation as to if and how said action will negatively affect sea turtles. (1)

### Species Status Completeness (Total Points: 5)

1. Does the consultation adequately describe the species and its habitat/critical habitat? (1)
2. Is the life history of the species addressed? (1)
3. Is there a detailed demographic analysis (if available for the species), including population size, variability and stability? (1)
4. Is the status and distribution of the species addressed, including reasons for listing? (1)
5. Is there an analysis of the species/critical habitat likely to be affected by the action? (1)

### Cumulative Effects Completeness (Total Points: 2)

1. Does the consultation consider the likelihood of the species to be able to recover? (1)
2. Does the consultation consider the effects of *future* State, tribal, local or private actions that are reasonably certain to occur, as per the handbook? (1)

## SI APPENDIX 2: SCORING RUBRIC FOR INFORMAL ESA SECTION 7 CONSULTATIONS

### Informal Criteria Baseline (Total Points: 5)

1. Mentions the action (1)
2. Some analysis of the action (1)
3. Some analysis of the impacted species (1)
4. Reason the consultation stayed informal is mentioned (1)
5. Map of the area affected by the action (1)

## SI APPENDIX 3: CONSULTATION PROCESS QUESTIONS FOR FISH AND WILDLIFE SERVICE AND NATIONAL MARINE FISHERIES SERVICE BIOLOGISTS

1. Can you tell me a bit about how the consultation process usually begins for you?
2. How frequently do you work on consultation? Has this number increased or decreased in recent years? Why might that be so?
3. How common is it to ask the action agency to provide more information on the action?
4. Have you seen a change over time in the way consultations are completed?
5. The number of consultations for FWS in Florida has been steadily decreasing since 2008 (according to the TAILS database there were 1099 in 2008 vs. 347 in 2014). Do you have an impression of how often you aren’t consulted on things?
6. Is there a consultation key for sea turtles, similar to the FWS Wood Stork Consultation Key? If not, is this something the Service would consider doing? Would this be an improvement to the process? Would you be in favor of a more standardized way to approach the consultation process? (Keys, a standardized ITP, etc.)
7. Can you explain the process of going through the literature and files on hand to satisfy the “best possible science” condition?
8. How do you exercise precaution when dealing with scientific uncertainty surrounding the effects of an action on a species/critical habitat? How much benefit of the doubt do you give to the species? Does it differ depending on the situation? Is this an issue you deal with on a regular basis?
9. How much time do you spend on the average consultation? FWS TAILS database says the average days for approval for formal consultations is 89 (13 for informal) days. Does that seem right?
10. Is pervious take ever tallied (formally or informally) to get a sense of how much has been done to a species over time? In your view, would this be a feasible/helpful thing to implement?
11. How often do you consult the section 7 Handbook?
12. Do you ever get requests for re-initiation of consultations?
13. NMFS is taking the lead on the revision of the handbook this year. What would you like to see in the revision? In your opinion, is there something that should be clarified?
14. What is your opinion on making all of the final documents publicly available (NMFS has PCTS, Vero Beach has the formal consultations online but not the informal documents)?
15. Where is there the most room for improvement in the consultation process? Does it work well as is?

## SI APPENDIX 4: BIOLOGIST RESPONSES

Included in Open Science Framework archive at https://dx.doi.org/10.17605/OSF.IO/KAJUQ

## SI APPENDIX 5: WOOD STORK CONSULTATION KEY

Included in Open Science Framework archive at https://dx.doi.org/10.17605/OSF.IO/KAJUQ

## SI APPENDIX 6: RECOVERY PLANNING AND IMPLEMENTATION FACT SHEET

Included in Open Science Framework archive at https://dx.doi.org/10.17605/OSF.IO/KAJUQ

## SI APPENDIX 7: SPECIES STATUS ASSESSMENT PRESENTATION

Included in Open Science Framework archive at https://dx.doi.org/10.17605/OSF.IO/KAJUQ

